# Mineralization kinetics during embryonic avian bone growth: a three-dimensional multiscale and cryogenic imaging approach

**DOI:** 10.64898/2026.04.13.718130

**Authors:** Anne Seewald, Jingxiao Zhong, Vedanti Sutaria, Issa Al Charkawi, Angelo Valleriani, Peter Fratzl, Emeline Raguin

## Abstract

Bone formation during embryonic development requires the rapid and sustained delivery of large amounts of calcium to mineralizing tissue. In avian embryos, this process coincides with a unique physiological transition in calcium supply, shifting from limited yolk reserves to massive mobilization from the eggshell. To address the question of how calcium transport responds to the increasing mineral demand during this transition, we combine micro computed tomography with three-dimensional cryogenic FIB-SEM imaging at different embryonic stages in the developing quail femur. Over this developmental period, bone mineralized volume increases rapidly through periosteal expansion. We observe that bone forming cells consistently contain numerous membrane-bound carriers loaded with mineral precursors throughout all growth stages. By integrating measurements across length scales, we calculate the intracellular velocities required to sustain the observed mineral deposition in the successive developmental stages. Surprisingly, these velocities remain within a similar range across development, despite the significant difference of bone formation rates, and are consistent with active transport by molecular motors. The increasing mineral demand corresponds to the expansion of mineralizing surfaces, which leads to an increasing number of cells, while the transport capacity of individual cells remains similar. Our work reveals a perfect tuning between the calcium transport capacity and bone growth, even in a situation where the skeletal growth is accelerating in the quail embryo.

**Significance statement:** Despite the importance of bone mineralization for the integrity of the skeleton, surprisingly little is known about how calcium is transported across the organic matrix to the mineralization sites during embryonal development. Using an avian model, where the main source of calcium shifts from a limited contribution from the yolk to the massive reservoir in the eggshell, we quantify bone mineral deposition using tissue level characterization together with three-dimensional cryogenic nanoscale imaging. We discover that intracellular vesicles containing mineral precursors are transported at similar speed, although the overall mineral deposition rate is increasing six-fold from embryonal day 10 to 14. This shows that the acceleration in mineralization activity is due to an increased number of bone cells rather than to a larger workload for individual cells. These findings demonstrate that rapid bone development is regulated to synchronize growth rates and mineral transport.

## 1. Introduction

Bone formation during embryonic development requires the continuous delivery of large amounts of calcium to sites of mineralization (1). In many vertebrate systems in which embryos develop within a continuously regulated physiological environment, calcium is continuously supplied through maternal circulation or dietary intake and maintained within a narrow physiological range by systemic homeostatic mechanisms. In contrast, in some oviparous systems such as birds, embryos develop within a closed system in which calcium availability undergoes a major transition, shifting from a limited yolk supply to the massive reservoir stored in the eggshell (2-4). This transition imposes a time-dependent constraint on skeletal mineralization, as calcium supply is progressively rearranged through systemic mobilization and transport mechanisms (5-8). Calcium must therefore be delivered to the developing skeleton despite a changing supply, raising the question of whether calcium transport adapts across development in magnitude and mode, or both.

In avian embryos, skeletal growth occurs over a short developmental period, generating substantial temporal variation in mineral demand. During early stages, calcium is supplied primarily from the yolk, where it is bound to highly phosphorylated proteins such as phosvitin (5). Progressive dephosphorylation of phosvitin mobilizes this limited reservoir and supports early skeletal formation (9). As development proceeds, the chorioallantoic membrane (CAM), a highly vascularized extraembryonic membrane responsible for gas exchange and shell resorption, progressively becomes the main route of calcium supply by dissolving the inner shell surface and transporting the released calcium into the embryonic circulation (2, 6, 8). This transition occurs over a defined developmental window during which the CAM establishes functional contact with the shell and increases its transport capacity, leading to a rapid rise in systemic calcium availability. This period coincides with the onset and acceleration of skeletal mineralization, during which both calcium supply and mineral demand increase in a synchronized manner (3, 7).

Bone formation involves both longitudinal growth through endochondral ossification (10-12) and radial expansion through periosteal apposition (13, 14). During radial expansion, rapidly deposited woven bone embeds a subset of osteoblasts that later differentiates into osteocytes (15). These newly formed osteocytes establish an interconnected lacuno-canalicular network that maintains communication with the surrounding matrix and contributes to the regulation of the mineralizing environment (16-18), expanding with bone growth and contributing to mineral transport within the tissue (16). As a result, skeletal growth is associated with a rapid increase in mineralized volume and therefore in total calcium demand of the developing bone.

To sustain this demand, calcium must be transported across multiple levels, from the circulation to the bone surface and ultimately to sites of mineral deposition within the tissue. At the cellular level, this involves regulated intracellular handling through the storage and trafficking of mineral precursors within membrane-bound structures observed in different species (1, 19-23). Intracellular compartmentalization limits exposure of the cytosol to high calcium concentrations and enables controlled delivery of mineral precursors to the extracellular matrix, enabling controlled mineral delivery while maintaining intracellular homeostasis (24). Consistent with this hierarchical organization, vesicles containing mineral precursors have been observed within the vasculature of the developing quail (25) and chicken (26) as well as in the murine growth plate (27), suggesting a coordinated transfer from the circulation to cells and ultimately to the extracellular matrix.

In our previous work using chick embryos (1), we showed that intracellular transport is an active process, with vesicles containing mineral precursors moving at velocities of molecular motor motion. However, during avian embryonic development, this transport operates within a changing physiological context where calcium availability increases as supply shifts from the yolk to the eggshell. Whether intracellular calcium transport adapts to this coupled increase in mineral demand and systemic supply change remains unknown. While the cellular and molecular regulation of bone formation has been extensively characterized (28, 29), quantitative integration of mineral demand with calcium transport across tissue and cellular scales during development remains limited. This limitation reflects the difficulty of resolving mineral precursors within their native three-dimensional cellular environment, where their distribution directly relates to intracellular transport. Recent advances in cryogenic volumetric imaging enable visualization of mineral precursors with their native cellular context (1, 30, 31). This provides direct quantification of mineral precursors and their relation to intracellular transport processes in the context of increasing mineral demand.

Here, we combine tissue level quantification of mineral growth using micro computed tomography (µCT) with cryogenic Focused Ion Beam-Scanning Electron Microscopy (cryo FIB-SEM) imaging of intracellular mineral precursors to determine how calcium delivery is sustained in the quail femur during development. By integrating mineral demand, vesicle cargo, vesicle density, and transport distances, we estimate the intracellular transport required at successive developmental stages. This multiscale approach allows us to determine whether rapid skeletal growth is supported by increased cellular transport capacity or by changes at the tissue level.

## 2. Results

### 2.1 Tissue level femoral growth is achieved by periosteal expansion and osteogenic proliferation

µCT imaging reveals rapid and accelerated growth of the quail femur from embryonic developmental day (EDD) 9 to 14 (Fig. 1) characterized by a nonlinear increase in mineralized volume over time. For each femur, a region of interest (ROI) corresponding to the central 15% of the total femoral length is examined to focus on diametric bone apposition and to ensure comparison of equivalent anatomical regions across developmental stages (Fig. 1A, orange). 3D cross-sectional views of the midshaft show progressive circumferential expansion through subperiosteal apposition of new mineralized trabeculae (Fig. 1B). Quantitative assessment reveals a strong and accelerated increase in median mineralized bone volume within the ROI, from 0.03 mm^3^ at EDD9 to 0.19 mm^3^ at EDD14 (Fig. 1C). This increase is well described by a quadratic fit (see Fig. 1C): the mineralized bone volume does not increase linearly but accelerates over developmental time. This non-linear increase corresponds to a three-dimensional mineralized bone formation rate of about 0.01 mm^3^/day at EDD10 that increases six-fold to 0,06 mm^3^/day at EDD14. In contrast, the bone volume fraction, defined as the ratio of the bone volume (BV) to the total volume of the region of interest (TV), increases only moderately from 0.26 to 0.30 (Fig. 1D) and follows an approximately linear trend. Thus, while total mineralized volume increases substantially, the relative proportion of mineralized tissue remains nearly constant. These results indicate that skeletal growth at the tissue level is achieved primarily through geometric expansion and apposition of new tissue, resulting in a rapid increase in the total mineral demand.

**Figure 1:**
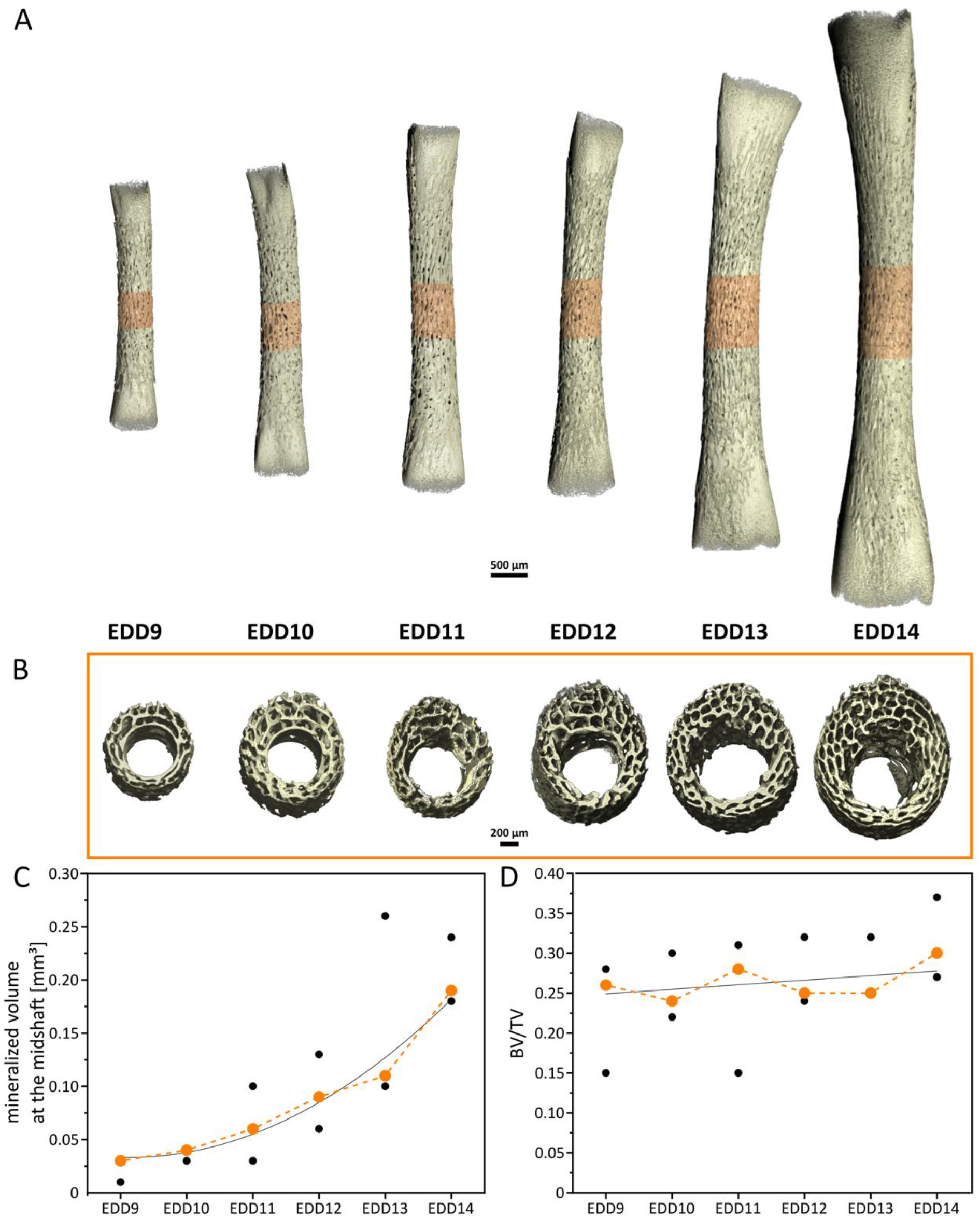
Development of the quail femur at different embryonic day (A) 3D orthographic projections of µCT reconstructions showing progressive femoral growth. The midshaft region of interest (ROI), defined as the central 15% of the total femoral length, is highlighted in orange (B) Cross-sectional 3D views of the midshaft (ROIs shown in A), illustrating trabecular apposition (C) Volume of mineralized bone at the midshaft ROI showing a quadratic fit (black line) of the median values (orange dot) highlighting the accelerated increase in mineralized volume across developmental stages (D) Bone volume fraction (BV/TV) showing a linear fit (black line) of the median value (orange dot) indicating a relatively constant proportion of mineralized tissue over time. Each data point (black dots) corresponds to a different embryo.

To identify the cellular processes supporting this radial expansion, osteogenic cell activity was assessed with histology and immunofluorescence. Fast Blue staining shows strong alkaline phosphatase (ALP) activity localized to the periosteal layer at all developmental stages (Fig. 2A), corresponding to active osteogenic and mineralization activity by osteoblasts. Immunofluorescence labelling for phosphorylated histone H3 (pHH3) reveals numerous proliferating cells concentrated in the outer periosteal region (Fig. 2B). These observations demonstrate that circumferential growth is supported by sustained osteogenic activity and expansion of the osteogenic cell population at the periosteal surface.

**Figure 2:**
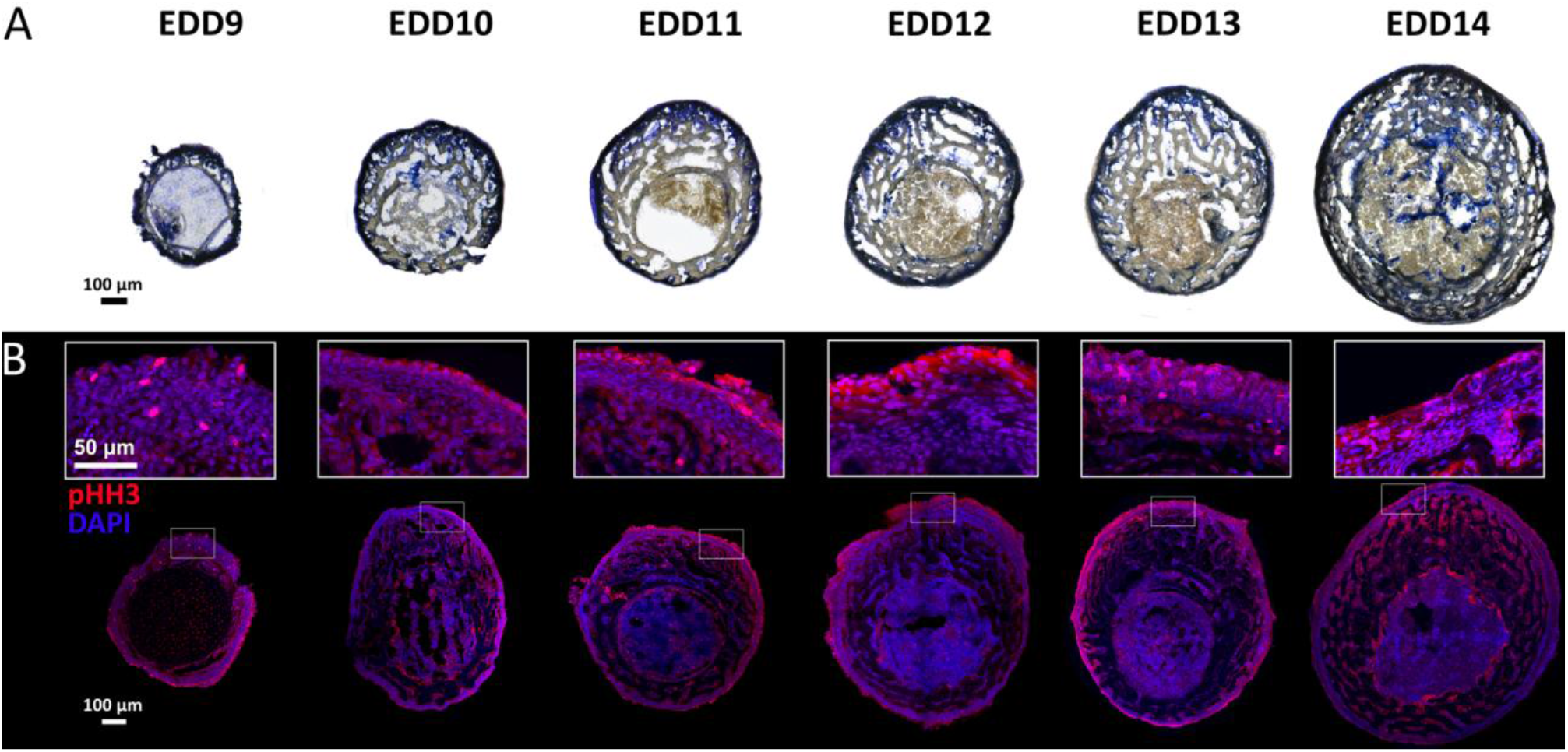
Osteogenic activity during femoral growth (A) Fast Blue staining for alkaline phosphatase (ALP) activity, visualized as blue precipitate, highlighting osteogenic activity at the periosteal surface. (B) Immunofluorescence staining for proliferating cells (phospho-histone H3, pHH3, in red) and nuclei (DAPI, in blue). The white box shows a higher-magnification view of the outer bone layer, highlighting the concentration of proliferating cells in the periosteal region.

### 2.2 Intracellular vesicular structures containing mineral precursors during embryonic development

Cryo FIB-SEM imaging of the mineralization front reveals numerous intracellular structures containing electron-dense mineral granules at all developmental stages examined (EDD9 to 12, Movie SI1 and Fig. SI2). These structures are observed in both osteocytes and cells located at the mineralizing surface, indicating that intracellular mineral carriers are a consistent feature of bone forming cells during skeletal embryonic development (Movie SI1 and Fig. SI2). In mixed InLens/secondary electron (SE) images, these vesicles are identified by their surrounding membrane (Fig. 3A and C, Movie SI1 and Fig. SI2A and C). Corresponding backscattered electron (BSE) images show bright particles within these carriers, confirming the presence of mineral precursors (Fig. 3B and D, Movie SI1 and Fig. SI2B and D). Three-dimensional rendering of the segmented structures confirms their intracellular localization and shows a high density of mineral carriers within the cells at the mineralization front (Movie SI1 and Fig. 3E). Some carriers display morphological features characteristic of mitochondria containing dense granules, whereas others exhibit a simpler vesicular morphology. Because mitochondria with dense granules have been reported to participate in intracellular mineral handling (22, 32, 33), they were included in the results. Here, the term “vesicles” refers to all intracellular membrane-bound structures containing electron-dense mineral cargo, potentially including mitochondria with dense granules. In addition, numerous vesicles and mitochondria without visible mineral granules were also observed.

**Figure 3:**
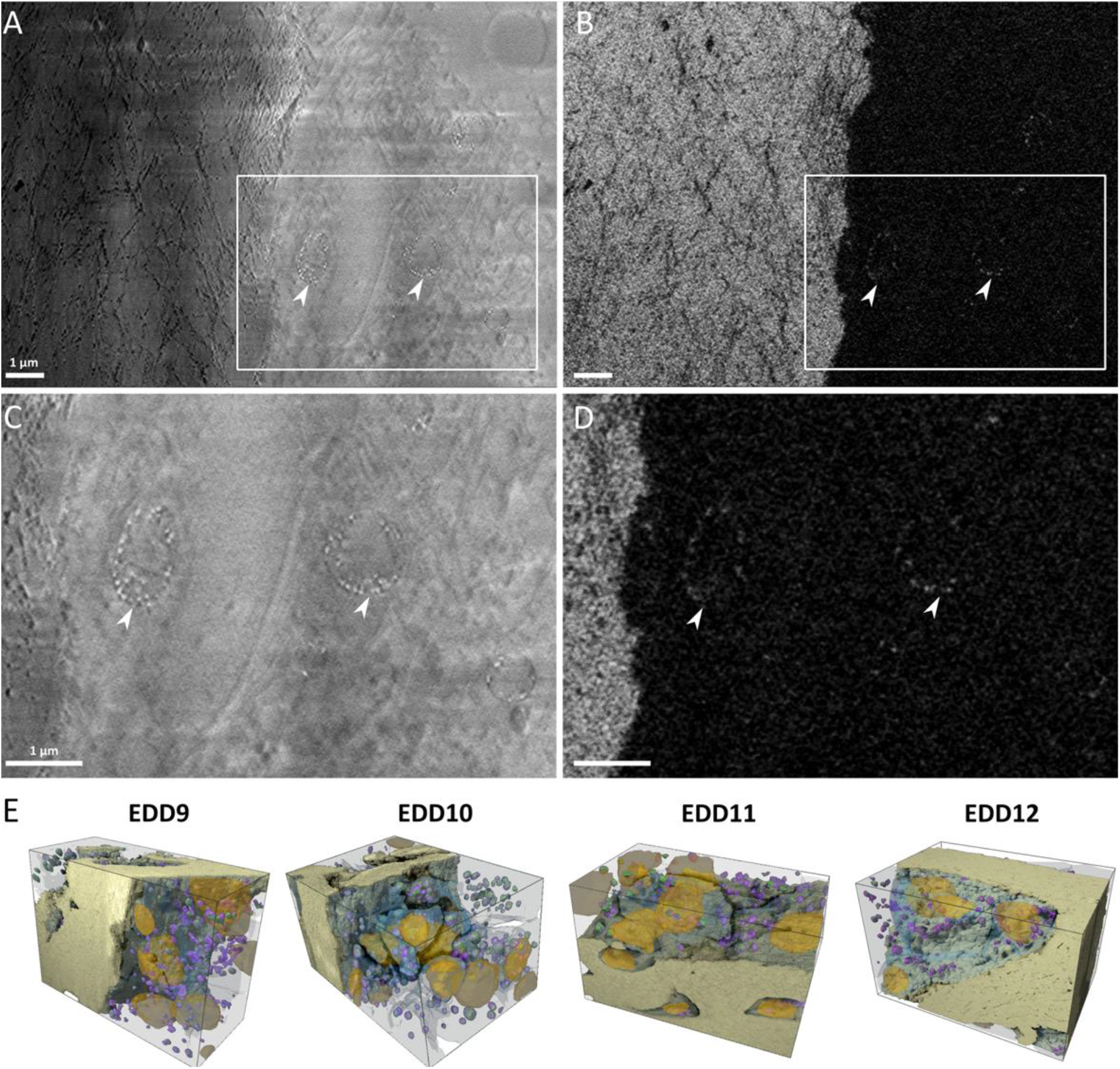
Representative images of intracellular vesicles containing mineral precursors. (A) Mixed InLens/SE image showing the presence of a continuous membrane (white arrows) at EDD10. (B) Corresponding backscattered electron (BSE) image of the same region, showing electron-dense particles within the membrane. (C) Higher-magnification view of the intracellular vesicles in mixed InLens/SE defined by the white box in (A). (D) Corresponding BSE image highlighting dense mineral precursors within vesicles. Comparative representative images of the different developmental stages are shown in Fig.SI2 (E) Three-dimensional perspective rendering of representative stacks at all the developmental stages investigated showing the spatial organization of intracellular vesicles. Segmented structures include bone (light yellow), cells (light blue/grey), nuclei (orange), vesicles (purple), and mineral precursors (green). All the vesicles containing mineral precursors are found within cells.

Quantitative analysis shows that the mineral precursors volume increases with vesicle size at all developmental stages (SI3A). Mineral precursor volume was plotted as a function of vesicle size on logarithmic scales, allowing a scaling exponent to be fitted, which reflects how mineral content varies with vesicles size and whether larger vesicles are proportionally more or less filled. A constant filling factor would correspond to an exponent of 1, where precursors and vesicle volume are just proportional to each other. An exponent lower than 1 then means that larger vesicles have a smaller filling factor. The fitted scaling exponents range from 0.63 (EDD11) to 1.07 (EDD12) (EDD9: 0.69; EDD10: 0.98), indicating that the relationship between vesicle size and mineral precursor volume changes across development rather than remaining proportional. The distribution of vesicle volumes is dominated by small carriers at all stages, with more than 90% of them smaller than 1 µm^3^ (SI3B). As a result, the fitted scaling exponents are influenced by the relatively small number of large vesicles and thus reflect the upper end of the size distribution rather than the dominant vesicle sizes. The vesicle filling factor, defined as the ratio between mineral volume and vesicle volume, provides a more representative measure of vesicle loading and varies across development (SI3C), with median values of 0.049 (EDD9), 0.066 (EDD10), 0.031 (EDD11), and 0.073 (EDD12). Interestingly, the median filling factor at EDD11 was approximately half of that measured at EDD12, revealing substantial heterogeneity in vesicle loading. When expressed as the contribution to the total precursor volume (SI3D), nearly 50% of the mineral is consistently packaged into vesicles smaller than 0.05 µm^3^ across all developmental stages. This shows that, despite variability in vesicle loading, mineral transport relies mainly on a large population of small carriers. Only at the latest developmental stage investigated here (EDD12), there is a relative change toward greater contribution from larger vesicles.

In addition to individual mineral carriers, larger membrane-bound compartments enclosing multiple vesicles are observed in a subset of cells (Fig. 4, Movie SI1 and Fig. SI4). These compartments show heterogeneous morphology, including spherical, ovoid, and elongated tubular shapes, and vary substantially in size. Measured volumes ranged from approximately 1.7 to 53.5 µm^3^, with the largest examples observed at EDD10 although some of these structures were only partially captured within the imaging volume. In mixed InLens/SE images, these compartments were delineated by a continuous membrane (Fig. 4A, Movie SI1 and Fig. SI4A and C), whereas BSE images showed that many of the internal vesicles contained electron-dense mineral precursors (Fig. 4B, Movie SI1 and Fig. SI4B and D).

**Figure 4:**
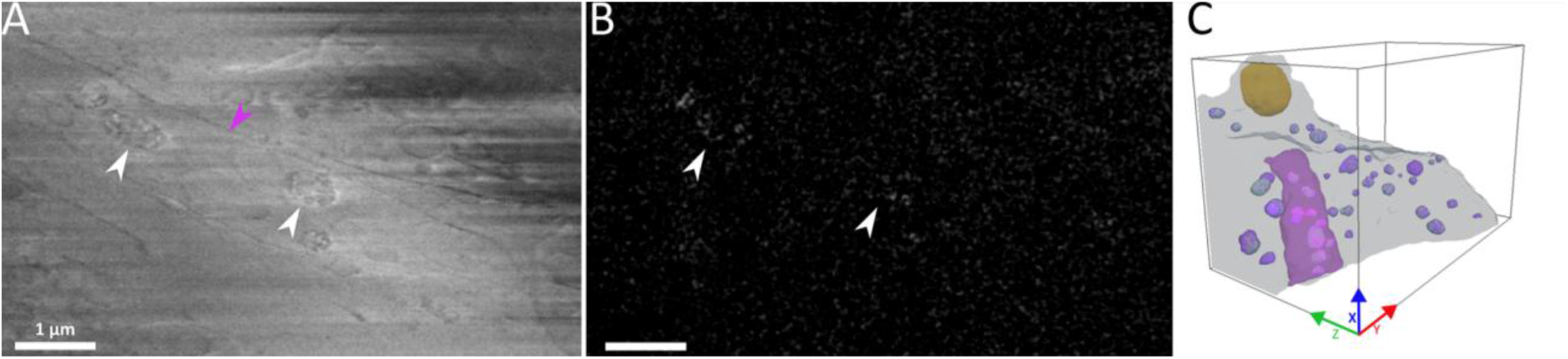
Representative images in the XZ plan of large membrane-bound structure containing vesicles filled with mineral precursors at EDD9 (A) Mixed InLens/SE image showing a large intracellular compartment delimited by a membrane (pink arrow) and containing multiple vesicles (white arrows) (B) Corresponding backscattered electron (BSE) image of the same region, revealing electron-dense particles within the internal vesicles (white arrows). (C) Three-dimensional perspective rendering illustrating the spatial organization of this membrane-bound compartment (pink) within the cell (light blue/grey) and the intracellular vesicles (purple) containing mineral precursors (green). Box dimension: X 11.52 µm, Y 17.34 µm, Z 10.33 µm. Comparative representative images of the different developmental stages where these structures are presents are presented in Fig. SI4.

Their occurrence varies markedly across developmental stages. A single example was detected at EDD9. At EDD10, they were present in all acquired stacks and occurred in relatively high numbers, including multiple compartments within the same cell in several cases. At EDD11, they were observed again in only one stack, whereas no such structure was visualized at EDD12. Within individual cells, these compartments were most often present as a single organelle, although up to six per cell were occasionally observed at EDD10. Three-dimensional segmentation rendering shows that each compartment contains multiple internal vesicles (Fig. 4C, Movie SI1 and Fig. SI4E), with counts ranging from 1 to 12 vesicles per compartment.

### 2.3 Kinetics and pathways of intracellular mineral transport during bone development

To relate tissue level mineral demand to intracellular transport, we extended an approach previously developed in chick embryos (1), in which vesicle transport is estimated from the amount of mineral delivered per cell, the intracellular mineral cargo, and the distance over which it must be transported. Here, this approach is applied at different developmental stages using directly measured volumetric parameters and statistical resampling. Briefly, the calculation integrates (i) the mineralized volume per cell derived from the µCT (SI5), (ii) the total volume of mineral precursors contained within intracellular vesicles, quantified from cryo FIB-SEM and extrapolated to a representative cell volume using linear regression and (iii) the intracellular transport distance defined as the path from the vasculature to the cell and approximated as half the trabecular thickness with half the distance between neighboring osteocytes. These parameters define a lower bound (*θ*) for vesicle velocity required to sustain mineralization (see Methods and SI5-6 for a detailed description of the calculation procedure and parameter distributions). Permutation resampling was used to generate distributions of required velocities, enabling statistical comparison between stages.

The estimated mean velocities are 0.7 ±0.3 µm/s at EDD9, 0.3 ±0.1 µm/s at EDD10, 1.1 ±0.5 µm/s at EDD11, and 0.6 ±0.2 µm/s at EDD12 (Fig. 5A). Despite the strong increase in mineralized volume during development, the distributions of estimated vesicle velocities overlap substantially and remain within the same order of magnitude. Values at EDD10 are lower and show reduced variability compared to the other stages. Both standard z-test and permutation testing did not detect significant differences between stages.

**Figure 5:**
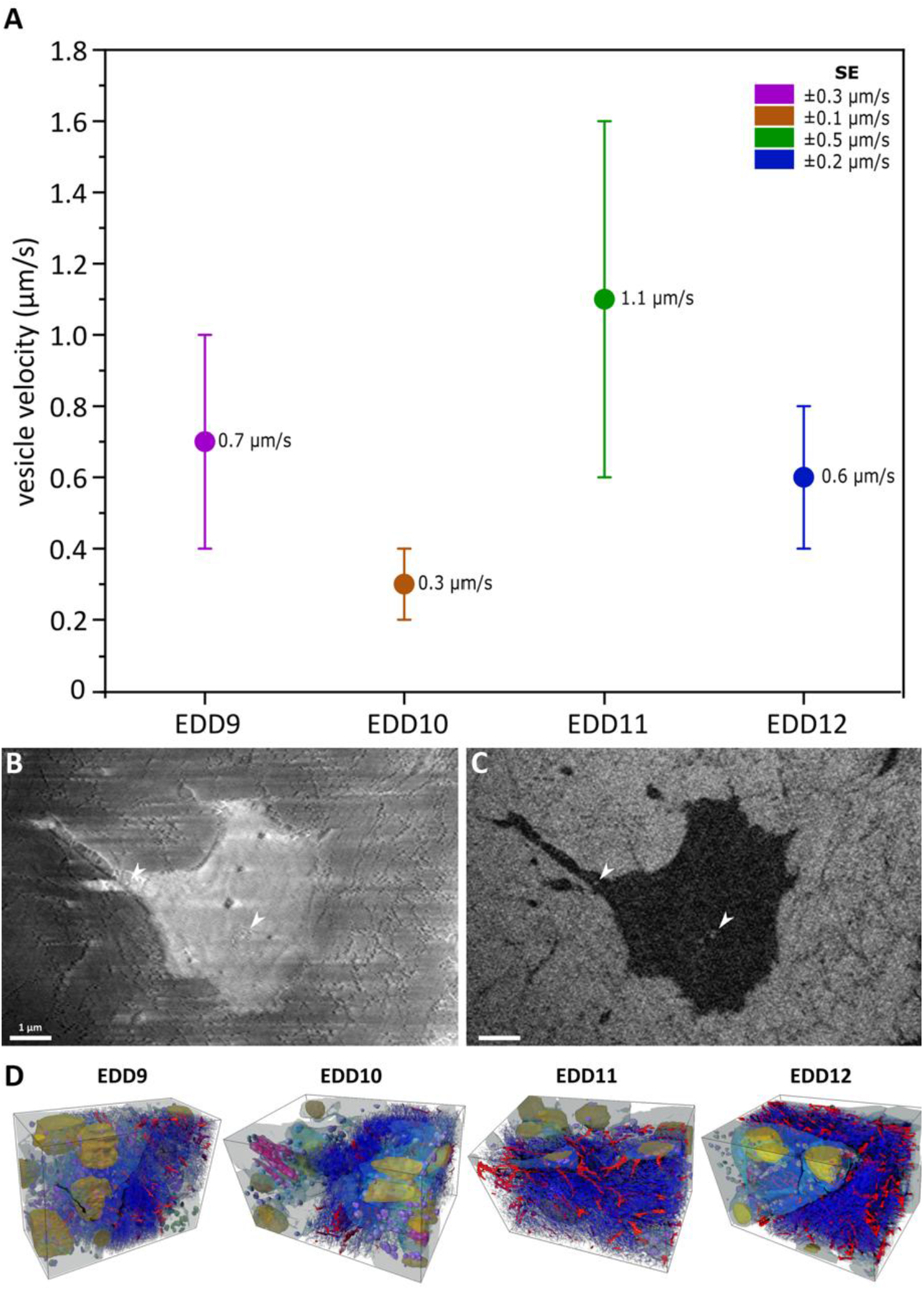
Intracellular velocities and transport routes in the mineralized bone (A) Graph representing the estimated lower bound of intracellular vesicle velocity required to sustain mineralization at each developmental stage, with their propagated error. (B) Mixed InLens/SE image at EDD10 showing an osteocyte and an extending canaliculus. The white arrow point to a vesicle at the entrance of the canaliculus and vesicles containing mineral precursors are seen within the cell (C) Corresponding backscattered electron (BSE) image of the same region, revealing the electron-dense particle within the osteocyte and one at the entrance of the canaliculus (white arrows). (D) Three-dimensional perspective rendering of the segmented lacuno-canalicular network and associated nanochannels illustrating their spatial organization. Segmented structures include canaliculi (red), nanochannels (blue), cells (light blue/grey), nuclei (orange), intracellular vesicles (purple) containing mineral precursors (green), and the large membrane-bound structure at EDD10 (pink).

For these precursors to contribute to matrix mineralization, pathways must exist for their transfer from cells and their distribution within the extracellular matrix. An electron-dense particle is observed within the lacuno-canalicular system at EDD10 (Fig. 5B, C). In backscattered electron images, this dense granule is detected at the entrance of a canaliculus extending from an osteocyte (Fig. 5C, white arrow). The corresponding mixed InLens/SE image allowed identification of the canalicular geometry, although the membrane contrast around the particle was less clearly defined. Within the same cells, intracellular vesicles containing mineral precursors were also present (Fig. 5B, C, white arrows). Three-dimensional segmentation of cryo FIB-SEM datasets highlights the spatial organization of osteocyte processes and their canaliculi, which connect the intracellular path to the surrounding matrix, together with a network of nanochannels distributed throughout the mineralized tissue (Movie SI1 and Fig. 5D). Quantitative analysis shows that the combined volume of canaliculi and nanochannels represents a similar fraction of the mineralized bone volume at each developmental stage, corresponding to approximately 10-13% of the mineralized bone volume. Nanochannels accounted for most of this space, whereas canaliculi contributed a smaller fraction.

### 2.4 Extracellular mineral deposition

Cryo FIB-SEM imaging shows extracellular mineral deposits within regions of newly formed matrix at all developmental stages examined (EDD9-EDD12) (Fig. 6). In mixed InLens/SE images (Fig. 6A, C), these regions are characterized by a loose and disorganized collagen network surrounding partially embedded cells. Corresponding BSE images (Fig. 6B, D) show extracellular deposits as bright foci (yellow arrows) distributed within these regions. The foci appear as discrete, irregularly shaped domains and are not associated with any surrounding membrane contrary to the vesicles containing mineral granules (Fig. 6, white arrows).

**Figure 6:**
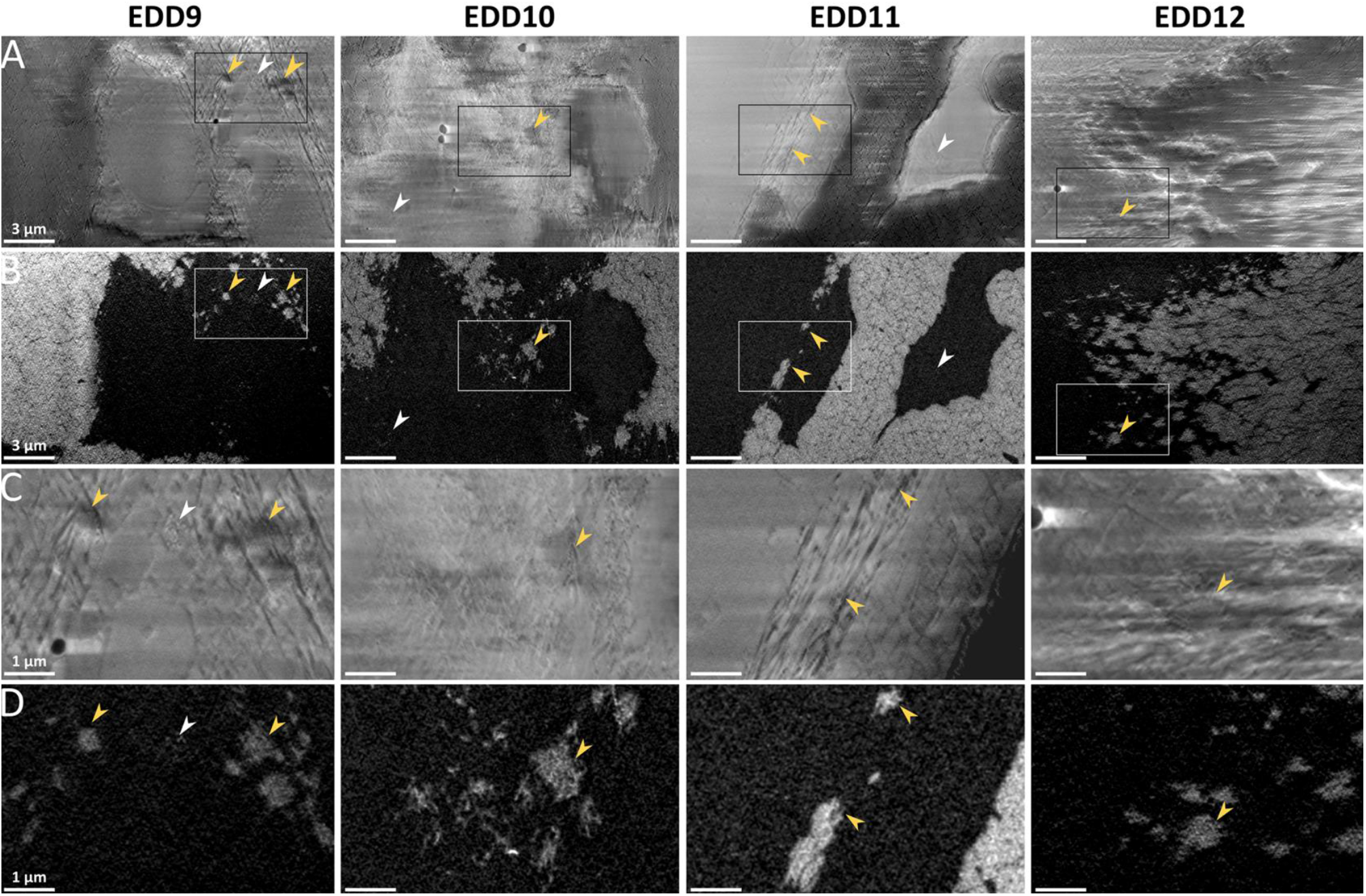
Extracellular mineral foci within unmineralized collagen fibrils observed from EDD9 to EDD12. (A) Mixed InLens/SE images showing disordered unmineralized collagen fibrils around pre-osteocytes at all developmental stages. (B) Backscattered electron (BSE) image of the same region as (A), highlighting denser mineral foci within the collagen fibrils. (C) Higher magnification of the disordered collagen shown in (A), with (D) being the corresponding images of (C) illustrating the denser mineral foci. Yellow arrows highlight denser foci within the collagen fibrils, whereas white arrows indicate dense mineral granules enclosed within intracellular vesicles.

## 3. Discussion

Skeletal development requires sustained delivery of large amounts of mineral to newly forming bone, a process that must remain effective despite simultaneous increases in mineral demand and changes in systemic calcium supply during avian development (4). Although vesicles containing mineral precursors have been described in bone forming cells, their role in supporting mineral delivery during development has remained largely descriptive (19-23). By combining tissue scale measurements of mineralized volume with 3D cryo FIB-SEM quantification of intracellular mineral carriers, this study provides a quantitative and dynamic interpretation of calcium transport during skeletal development. Extending a previously established multiscale approach in the chicken embryo (1), we evaluate intracellular transport requirements across successive developmental stages in Japanese quail where both mineralized tissue and systemic calcium supply increase in parallel as the source shifts from yolk to eggshell with the CAM. Our results show that intracellular calcium transport operates in a steady regime across development, and that the increasing mineral demand is accommodated by expansion of the mineralizing surface and recruitment of additional osteogenic cells, rather than by changes in vesicle transport dynamics.

The developmental window investigated (EDD9 to 12) corresponds to a period of major physiological reorganization in calcium availability. In chicken embryos, calcium contribution from the yolk decreases during early incubation and reaches a plateau around EDD11-12, at which point the eggshell becomes the dominant mineral source as the CAM becomes fully functional and continues to expand its transport capacity (3, 34, 35). Although the CAM is in contact with the shell starting at EDD9, effective calcium mobilization begins only after differentiation of specialized chorionic cells responsible for shell dissolution (36, 37). Comparative studies indicate that this transition occurs slightly earlier in Japanese quail, with EDD9 and EDD11-12 in chicken corresponding to EDD8-9 and EDD9.5-10 in quail respectively (38). The stages examined here therefore span this critical transition in calcium supply, during which systemic availability is reorganized while mineralized volume increases rapidly. Importantly, the present study does not directly measure calcium availability but instead relates intracellular organization to this known physiological transition.

The velocities calculated here remain of the same order of magnitude across developmental stages, indicating that intracellular trafficking operates within a stable dynamic regime despite progressive shifts in calcium supply. A distinct deviation is observed at EDD10, where both mean velocity and variability decrease, followed at EDD11 by higher values, although these differences are not statistically significant. This stage may therefore represent an adaptation phase in which calcium supply and mineral demand are not yet fully matched, consistent with the onset of eggshell calcium input, whereas EDD11 reflects a stage where this supply is fully active while mineralized volume continues to increase. The velocities measured in quail are higher than those previously estimated in chicken embryos at EDD13 (∼0.3 µm/s) (1), although they remain within the same order of magnitude. One possible explanation is that the present calculation incorporates a refined formulation based on directly measured volumetric parameters and propagated variability, which may lead to higher estimates compared to the earlier calculations. Alternatively, the faster developmental pace of quail may require comparable mineralization events to occur over shorter timescales, resulting in higher vesicle velocities. This supports the idea that calcium transport remains constrained within a biologically similar range across avian species, while the exact values shift with developmental timing and mineral demand.

The magnitude of these velocities is characteristic of active transport with molecular motors along microtubules. Reported velocities of kinesin and dynein (0.3 to 1 µm/s) (39-41) closely match the range measured here. Although vesicle loading varies at the level of individual carriers, the overall transport remains dominated by a large population of small vesicles at all stages, resulting in a conserved transport regime. At all stages, vesicles exhibit heterogeneous mineral load, and although mineral cargo increases with vesicle size, this relationship varies across development rather than following a consistent scaling behavior. However, when considering the contribution to the total mineral precursor volume, a consistent pattern emerges: across all stages, a substantial fraction of the mineral is carried by small vesicles. Differences occur within the developmental window corresponding to the transition in systemic calcium supply, suggesting that intracellular calcium handling responds to changes in calcium availability rather than altering transport dynamics. Previous structural studies in bone and cartilage cells have shown that mineral precursors are packaged within intracellular vesicles and that their loading is controlled by cellular ion homeostasis and intracellular vesicular storage, including lysosomal processing, mitochondria and endoplasmic reticulum (42, 43). The differences observed in vesicle filling between the embryonic stages may therefore reflect changes in how calcium is handled and packaged within cells, rather than changes in transport velocity itself. In addition, experimental and theoretical studies have shown that although cargo load influence vesicle motion when transport capacity is limited, intracellular velocity is determined mainly by the number of engaged molecular motors and by the organization of the cytoskeletal network (44, 45). However, changes in cargo amount or composition affect run length, persistence, or delivery efficiency rather than velocity itself (44).

Beyond individual mineral carriers, larger membrane-bound compartments enclosing several of these vesicles were predominantly observed at EDD10 and were absent at EDD12. These compartments show a wide range of sizes (1.7 to 53.5 µm^3^) and contained multiple vesicles below 1 µm in diameter. They are consistent with multivesicular structures of the endosomal and lysosomal system involved in sorting, cargo storage, and regulated secretion (24, 46). Although here, these structures are larger than classical multivesicular bodies described in many cell types, enlarged endo-lysosomal compartments have been reported in highly active secretory and mineralizing cells (24, 33, 47). Their preferential occurrence at EDD10 further supports the idea that this stage represents a transitional phase of intracellular buffering during the change in calcium supply. Around EDD10 in quail, calcium input from the eggshell through the CAM becomes established and progressively increases (2, 7) while larger amounts of calcium are incorporated into the expanding skeletal tissue. These processes appear synchronized during development. These compartments may contribute to short-term intracellular buffering of mineral, smoothing local variations as systemic supply and skeletal demand become progressively aligned during the yolk/eggshell transition. If these structures act as transient storage site, the vesicles they contain would not directly contribute to ongoing mineral delivery but excluding them in the kinetic estimates provides similar results, likely because they represent a very small portion relative to the total population of carriers. Their precise identity and function remain to be established.

At the tissue level, the increase in mineralized volume follows an accelerated, nonlinear trajectory, indicating that mineral demand rises disproportionately during later developmental stages and is associated with geometric enlargement of the developing cortical bone and with positive ALP staining localized at the periosteal surface. The periosteum serves as a major source of osteoprogenitor cells, and cortical growth proceeds through their recruitment, proliferation, and differentiation (48). Consistent with this mechanism, the ALP region observed here coincides with proliferating cells identified by pHH3 staining, indicating that new mineralizing fronts are established through expansion of the osteogenic population at the bone surface. As mineral deposition is localized to this peripheral region, bone enlargement expands the mineralizing surface, contributing to the increased mineralized volume. ALP activity promotes mineral formation by increasing inorganic phosphate availability and reducing inhibitors such as pyrophosphate (49). Thus, local calcium delivery becomes a key factor controlling mineral nucleation and the spatial distribution of mineral deposition. The discrete extracellular mineral foci observed at all stages likely correspond to nucleation events occurring where calcium delivered by actively transported intracellular vesicles is released into a phosphate rich extracellular environment favorable for mineral precipitation and crystal growth.

The structural and kinetic variations observed during embryonic skeletal growth indicate that the transition from the yolk to the establishment of the CAM as the dominant source of calcium most likely occurs between EDD10 and EDD11, despite staging comparisons with the chicken showing an earlier onset between EDD9 and EDD10 (38). Quail embryonic skeletal development is achieved while synchronized changes in calcium supply and skeletal growth occur, with intracellular calcium handling and tissue level expansion allowing efficient mineralization, rather than by increasing the transport capacity of individual cells. Calcium delivery follows the hierarchical pathway previously hypothesized (1) from vascular supply (25) to intracellular vesicular active transport and subsequent distribution through the lacuna-canalicular network.

## 4. Material and methods

### Eggs procurement and incubation

No approval by an ethics committee for animal experimentation was required for this study in accordance with the German Animal Welfare Act and the Laboratory Animal Welfare Ordinance. Fertilized Japanese quail (*Coturnix coturnix japonica*) eggs were obtained from a commercial breeder (Vogelsberger Wachtelzucht GmbH, Mücke, Germany and Wachtel-Shop Michael Volk e.K., Obersulm, Germany). Upon delivery, the eggshells were gently disinfected by wiping with 70% ethanol to reduce microbial contamination and were subsequently transferred to a digitally controlled egg incubator (BRUJA Flächenbrüter 3333D, Brutmaschinen-Janeschitz GmbH, Hammelburg, GE). Incubation was carried out at 38.5 ± 0.5 °C and 55 ± 5% relative humidity, with automated hourly turning. Embryos were euthanized via cervical dislocation and femurs were immediately dissected at embryonic developmental days (EDD) 9 to 14.

### Micro-Computed Tomography (µCT)

µCT was performed using an EasyTom 150/160 system (RX Solutions, Chavanod, France). For each embryonic stage (EDD9 to EDD14), three complete femurs were mounted individually in pipette tips filled with 70% ethanol. Imaging was carried out using the nano-focus tube (160) combined with a flat panel detector. Scans were acquired at 70 kV and a tube current of 80 µA, with a voxel size of 1.8 µm. Volumetric datasets were reconstructed from the acquired two-dimensional projections using X-Act software (RX Solutions, Chavanod, France). The resulting 3D volumes were saved as TIFF image for subsequent image processing and quantitative analysis.

### Image processing, segmentation, and quantitative assessment of µCT data

Mineralized bone volume was quantified at the femoral midshaft using Dragonfly software (Object Research Systems (ORS) Inc, Montreal, Canada). For each femur, the total bone length was first determined, and a midshaft region corresponding to 15% of this length was defined around the middle of the bone. For each image slice, the grayscale intensity histogram was analyzed to determine the transition between mineralized and non-mineralized phases by identifying the point at which the second derivative of the histogram approached zero, corresponding to a plateau in intensity values. The second-derivative evaluation was implemented using a custom Python script (Version 3.8.8). A global segmentation threshold was then defined as the mean of these slice-specific values across the volume of interest, and the associated standard deviation was used to estimate threshold uncertainty. Mineralized bone volume was subsequently computed from the resulting threshold-based segmentation.

Osteocyte lacunae were segmented using the Segmentation Wizard in Dragonfly ORS (Object Research Systems (ORS) Inc., Montreal, Canada). A supervised deep-learning approach based on a 2.5D U-Net model was employed, using approximately six manually training images to distinguish lacunae from the surrounding mineralized matrix and background. The resulting model was applied to all midshaft volumes, and a watershed-based separation method was employed to individualize adjacent or partially connected lacunae prior to quantitative assessment.

The average distance between osteocytes was quantified using a nearest-neighbor approach. After segmentation and watershed separation, each lacuna was considered as an individual object and its three-dimensional center of mass was computed in Dragonfly ORS (Object Research Systems (ORS) Inc., Montreal, Canada). The centroid coordinates (X, Y, Z) were obtained in physical units (µm) and exported for each volume. Nearest-neighbor distances were then calculated using a custom Python script in the Dragonfly embedded Anaconda Python 3 environment. For each osteocyte, the Euclidean distance to all other osteocyte centroids in three-dimensional space was computed, and the smallest value was defined as the nearest-neighbor distance.

To estimate the theoretical bone volume associated with individual osteocytes, a Voronoi spatial partitioning approach was applied. This methodology relies on the assumption that each osteocyte primarily influences mineralization within its local vicinity. Accordingly, the surrounding mineralized matrix was subdivided into discrete regions based on proximity to osteocyte lacunae, following the principles of a Voronoi tessellation: each bone voxel was assigned to the nearest osteocyte with the Euclidean distance principle, ensuring contiguous and non-overlapping spatial domains. Based on the voxel to cell attribution, distance maps were generated to show the mineralized bone voxels to their corresponding osteocytes. The total bone volume attributed to each osteocyte was then calculated by summing the voxels within its associated region. All Voronoi computations were implemented using a custom MATLAB script (R2021a, The MathWorks, Inc.). Fig. SI5A illustrates this approach with a representative distance maps and osteocyte lacunae-associated bone volumes visualized by color coding. The registered mineralized bone volume per osteocyte (*V*_*m*_) and the anatomical position of each cell were then evaluated spatially. Each cell’s position was expressed in cylindrical coordinates (*r, θ, z*) with respect to the central longitudinal axis of the femur. For each sample, the radial dimension was divided into 36 bins, and for each radial band, the regional average mineralized bone volume per cell (*V*_*m*_), the change in bone volume across the radius bands (Δ*V*_*m*_), and the number of cells detected within that band were calculated. Radial profiles of Δ*V*_*m*_ and osteocyte number were evaluated to assess the reliability of *V*_*m*_ estimates across the femoral cross-section (Fig. SI5B).

Trabecular thickness was quantified from µCT datasets using Dragonfly ORS (Object Research Systems (ORS) Inc., Montreal, Canada). Quantitative assessment was restricted to the actively mineralizing region by defining a region of interest encompassing only the growing bone front. The segmented mineralized bone volume was used as a mask to compute a volume thickness map, which provides the local thickness defined as the diameter of the largest sphere that fits entirely within the object at that location. Mean trabecular thickness was then obtained by averaging the thickness values across all voxels within the masked region of interest.

### Cryo-embedding and immunofluorescence staining

Immediately after dissection, femurs from embryonic developmental stages 9 to14 were cryo-embedded following dehydration in ascending sucrose solutions (10%, 20%, and 30% in distilled water; 24h each at 4°C). For embedding, a metal mold was precooled in isopropanol chilled with liquid nitrogen and filled with Tissue-Tek® O.C.T.™ Compound. Samples were positioned centrally within the mold, covered with embedding medium, and frozen while avoiding direct contact between the specimen and the precooled isopropanol. Cryosections of 20 µm thickness were prepared using a cryostat (Leica CM3060S, Leica Microsystems CMS GmbH, Wetzlar, Germany). For immunofluorescence staining, sections were blocked for 1 h at room temperature in blocking buffer (1% bovine serum albumin [BSA], 0.1% Tween-20, 0.1% dimethyl sulfoxide [DMSO] in PBS). Primary antibodies, 4’,6-diamidino-2-phenylindole DAPI (1:1000, Roche) and anti-pHH3 (1:100, 06-570, Sigma-Aldrich Chemie GmbH) were diluted in blocking buffer and incubated overnight at 4 °C. After washing (0.1% Tween-20, 0.1% DMSO in PBS), secondary antibodies (Alexa Fluor 555, 1:1000, A-21429, Abcam) were applied for 1 h at room temperature. Slides were mounted using Dako fluorescence mounting medium (S302380-2, Agilent Technologies, Santa Clara, USA).

Imaging was performed using confocal laser scanning microscopy (CLSM; Leica TCS SP8 DLS, Multiphoton, Leica Microsystems CMS GmbH, Wetzlar, Germany). A 20× HC PL APO CS2 dry objective (NA 0.75) was used for overview imaging and a 63× HC PL APO CS2 oil objective (NA 1.4) was used for high-resolution imaging of the mineralization front. DAPI fluorescence was excited with a 405 nm diode laser (λ_ex = 405 nm; λ_em = 415-485 nm), and Alexa Fluor 555 with a 561 nm DPSS laser (λ_ex= 561 nm; λ_em = 565-620 nm). Images were acquired as z-stacks with a resolution of 1024 × 1024 pixels (pixel size = 284 nm) at a scan speed of 600 Hz with consistent line averaging. Tile scans were performed when necessary to cover the full region of interest.

### Fast Blue staining for Alkaline Phosphatase (ALP)

Cryosections of the femoral bone from EDD9 to EDD14 were stained for alkaline phosphatase (ALP) activity using a freshly prepared Fast Blue reaction solution. The staining solution consisted of 0.9% NaCl, naphthol AS-MX (2.5 mg/mL), borax (5%, 750 µL), MgSO_4_ (10%, 40 µL), and Fast Blue RR salt (10 mg/mL). The solution was adjusted to pH≥ 8.5 using pH paper and filtered before application. Sections were air-dried for 30 min, followed by washing in 1× PBS and 0.9% NaCl (5 min each). Staining was performed by incubating sections with 200 µL of the reaction mixture for 15 min at room temperature in the dark, followed by two washes in PBS and a final wash in 0.9% NaCl. Stained sections were imaged using a Keyence Digital Microscope (VKX-5550E, Keyence, Germany) at 1000× magnification.

### Cryo Focused Ion Beam-Scanning Electron Microscopy (FIB-SEM)

Within minutes after sacrifice, right femoral cross-sections (∼400 µm thick) were cut from the midshaft of quail embryos from EDD9 to EDD12 using a double blade guillotine. Each bone section was immediately placed between gold-coated copper type B freezer hats, or a combination of one type B and one type A with 100 µm cavity (BALTIC preparation, Wetter, Germany). A 20 wt.% dextran solution (Sigma, 31390) was added as cryoprotectant. The freezer hats were arranged to form a cavity with a total height of 0.4 to 0.6 mm. Samples were cryo-immobilized using a Leica ICE high-pressure freezing (Leica Microsystems, Vienna, Austria). Frozen sample carriers were mounted onto a cryogenic sample holder in the Leica EM VCM loading station under liquid nitrogen and transferred via the VCT500 shuttle to a Leica ACE600 system (Leica Microsystems, Vienna, Austria) for freeze fracture and sputter coating. After fracturing the sample, a 10nm layer of carbon was sputter-coated onto the exposed surface, followed by a 10nm layer of platinum. Finally, the samples were transferred to the Zeiss Crossbeam 540 (Zeiss Microscopy GmbH, Oberkochen, Germany) using the VCT500 shuttle. Throughout the entire process, the samples were kept at a temperature below −145 °C.

### Serial Surface View (SSV) imaging

FIB-SEM serial surface view imaging was performed on three femoral midshaft sections per developmental stage using a Zeiss Crossbeam 540 dual-beam microscope (Zeiss Microscopy GmbH, Oberkochen, Germany). Samples were elevated to a height of 5.2 mm to align with the coincident point of the electron and ion beams and tilted at an angle of 54°. A trench measuring approximately 40 µm in length and 70 µm in width was milled using an ion beam current ranging from 1.5 to 30 nA. The newly exposed surface was subsequently polished and imaged using reduced ion beam currents (700 pA to 1.5 nA). Electron imaging was performed at an accelerating voltage of 2 keV and a probe current of 90 pA for all image stacks.

Images were sequentially acquired with an in-plane pixel size of 8 nm (x and y) and a slice thickness (z) of 8 nm to keep isotropic voxel dimensions. Each image stack was acquired at a resolution of 4096 × 3072 pixels in 8-bit grayscale using dual-channel acquisition mode, with simultaneous collection from the mixed InLens/secondary electron (SE) detector and the backscattered electron (BSE) detector.

### Image processing and segmentation

FIB-SEM image stacks were aligned, processed, and segmented using Dragonfly software, Version 2024 and 2025 (Object Research Systems (ORS) Inc., Montreal, Canada). The mixed InLens/SE images were first automatically aligned using the sum of square differences (SSD) registration algorithm, with manual fine adjustments applied when necessary. The resulting transformation parameters were subsequently applied to the corresponding backscattered electron (BSE) image stacks. Curtaining artifacts resulting from ion beam milling were corrected using a vertical destriping filter. To improve structural visibility, 3D contrast enhancement and noise reduction were applied using convolution filter. In addition, contrast in the mixed InLens/SE image stacks was further optimized using contrast-limited adaptive histogram equalization (CLAHE).

Segmentation of high-density, mineralized regions in the BSE image stacks was performed with the Segmentation Wizard module in Dragonfly, using a 2.5D U-Net deep learning model. The model was trained on five to seven manually segmented representative slices per dataset with variable input size. The trained model was subsequently applied to the entire image stack. The resulting segmentation was further refined by manual correction using the brush tool. In addition, a separate Segmentation Wizard model was trained and applied to the BSE image stacks to segment nanochannels and canaliculi. Cellular structures, including vesicles, nuclei, and cells, were segmented from the mixed InLens/SE image stacks using the active contour segmentation method available in the software.

### Kinetic estimation and statistical analysis

The intracellular velocity of vesicles carrying mineral precursors was estimated using a volumetric transport model adapted from our previous work in the chick embryos (1) and reformulated here using directly measured parameters in the different developmental stages. The required transport velocity was expressed as a lower bound (θ), defined as:

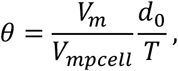

Where *V*_*m*_ is the mineralized volume of bone attributed to a cell over a period *T* (one day), *V*_*mpcell*_ is the total volume of mineral precursors contained within vesicles in a representative bone cell, and *d*_0_ is the intracellular transport distance. The mineralized volume per cell (*V*_*m*_) is obtained from µCT using the Voronoi approach as previously described (SI5). The transport distance (*d*_0_) is also obtained from µCT measurements as the sum of half the trabecular thickness and half the average intercellular distance within the midshaft region, representing the maximal intracellular path from vascular calcium supply to the mineralization site. *V*_*mpcell*_ corresponds to the vesicle mineral volume obtained from the segmented volume in cryo FIB-SEM. Because imaged cells were not captured entirely, values were extrapolated to a representative bone cell volume of 1000 µm^3^ using linear regression between cell volume and vesicular mineral content. Velocity estimates were computed independently for each developmental stage (EDD9 to EDD12) using three different samples for µCT and three independent datasets also from different samples for cryo FIB-SEM.

For each EDD, we have first computed the mean 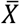of the product *V*_*m*_*d*_0_ out of the three independent samples and the mean 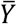of the three independently extrapolated *V*_*mpcell*_. Secondly, we have computed the standard errors 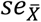and 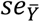after demonstrating that the variance across the biological samples is independent of the EDD. Therefore, the standard errors were computed using all 12 samples for 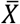and all 12 samples for 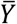. Practically, this means that those standard errors are the same for all EDD. Finally, for each EDD we have computed the corresponding velocity *θ* and its associated standard error *se*_*θ*_ using the method of error propagation, leading to:

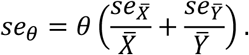

Statistical comparisons between developmental stages were performed using pairwise t-test and permutation significance testing. All measured data, intermediate values, calculation procedures and statistical model are provided in the Supplementary Information (SI6).

## Acknowledgements

We thank Ernesto Scoppola (MPIKG) for his assistance with figure visualization, and Jeannette Steffen (MPIKG) for her support with histological sample preparation and immunostaining.

