## Supplementary information for "Mineralization kinetics during embryonic avian bone growth: a three-dimensional multiscale and cryogenic imaging approach"

**Movie S1 (separate file).** Three-dimensional datasets of representative volumes acquired by cryo FIB-SEM are shown for the successive developmental stages investigated (EDD9-12). Volumes are displayed using the InLens/SE signal and the backscattered detector and the segmentation highlights the cells, nuclei, intracellular vesicles containing mineral precursors, enlarged compartment, canaliculi and nanochannels.

**Figures**

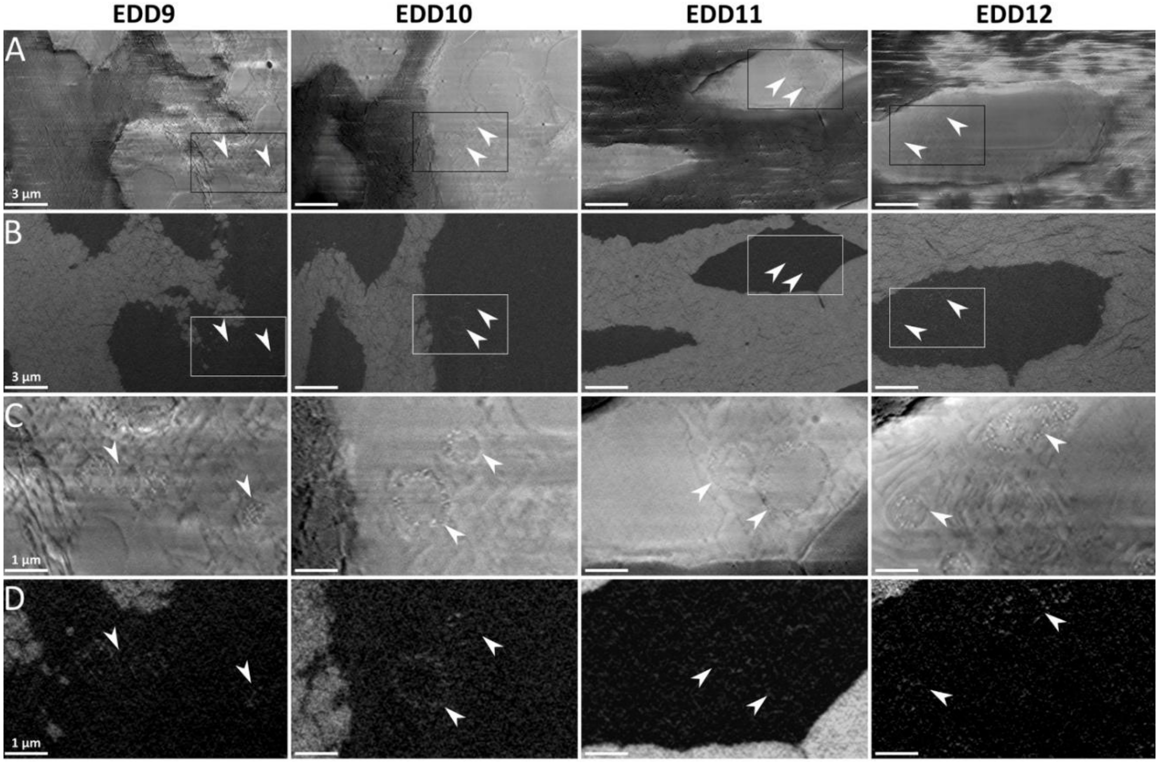


**Fig. S2.** Representative images of intracellular vesicles containing mineral precursors in all the stages investigated (A) Mixed InLens/SE image showing the presence of a continuous membrane (white arrows) (B) Corresponding backscattered electron (BSE) image of the same region, showing electron-dense particles within the membrane. (C) Higher-magnification view of the intracellular vesicles in mixed InLens/SE defined by the white box in (A). (D) Corresponding BSE image highlighting dense mineral precursors within vesicles.

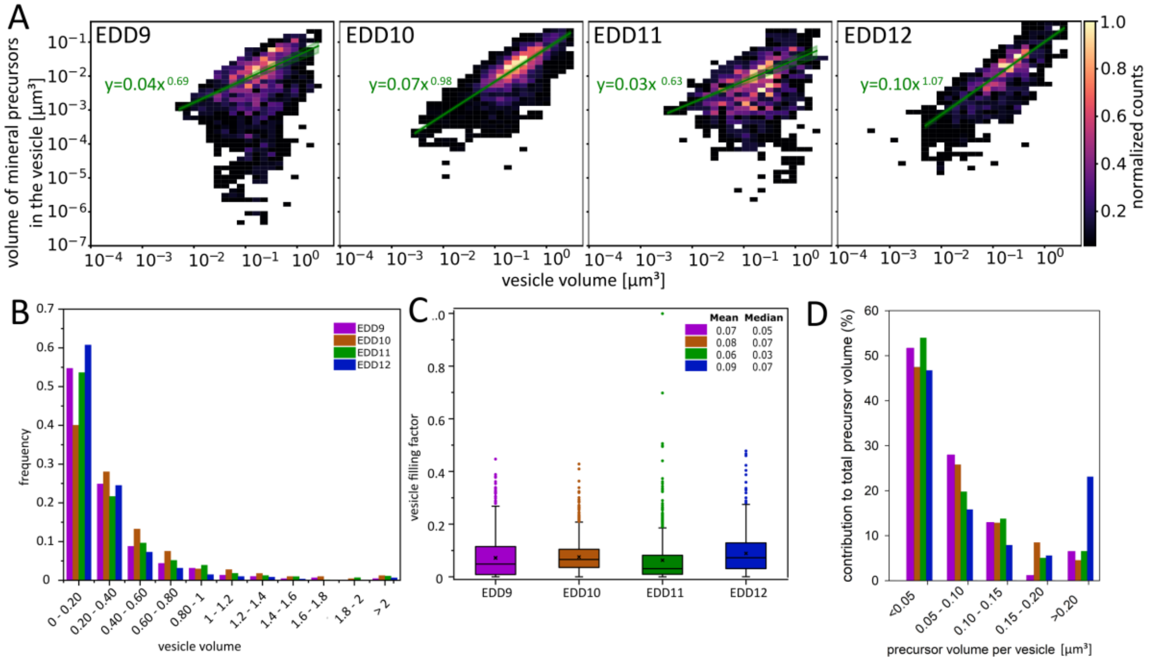

**Fig. S3.** Quantitative analysis of vesicles containing mineral precursors at all developmental stages investigated. (A) Density plot of mineral precursor volume in function of the vesicle volume, displayed on a double logarithmic scale. Color intensity reflects the local density of data points, with brighter regions indicating higher concentration. A fitted regression line (green) with 95% confidence band is overlaid, along with the corresponding equation. (B) Histogram showing the relative frequency of vesicles grouped by volume. (C) Filling factor (ratio of mineral precursor volume to vesicle volume) plotted per stage. (D) Histogram showing the contribution of each vesicle size class to the total mineral precursor volume.

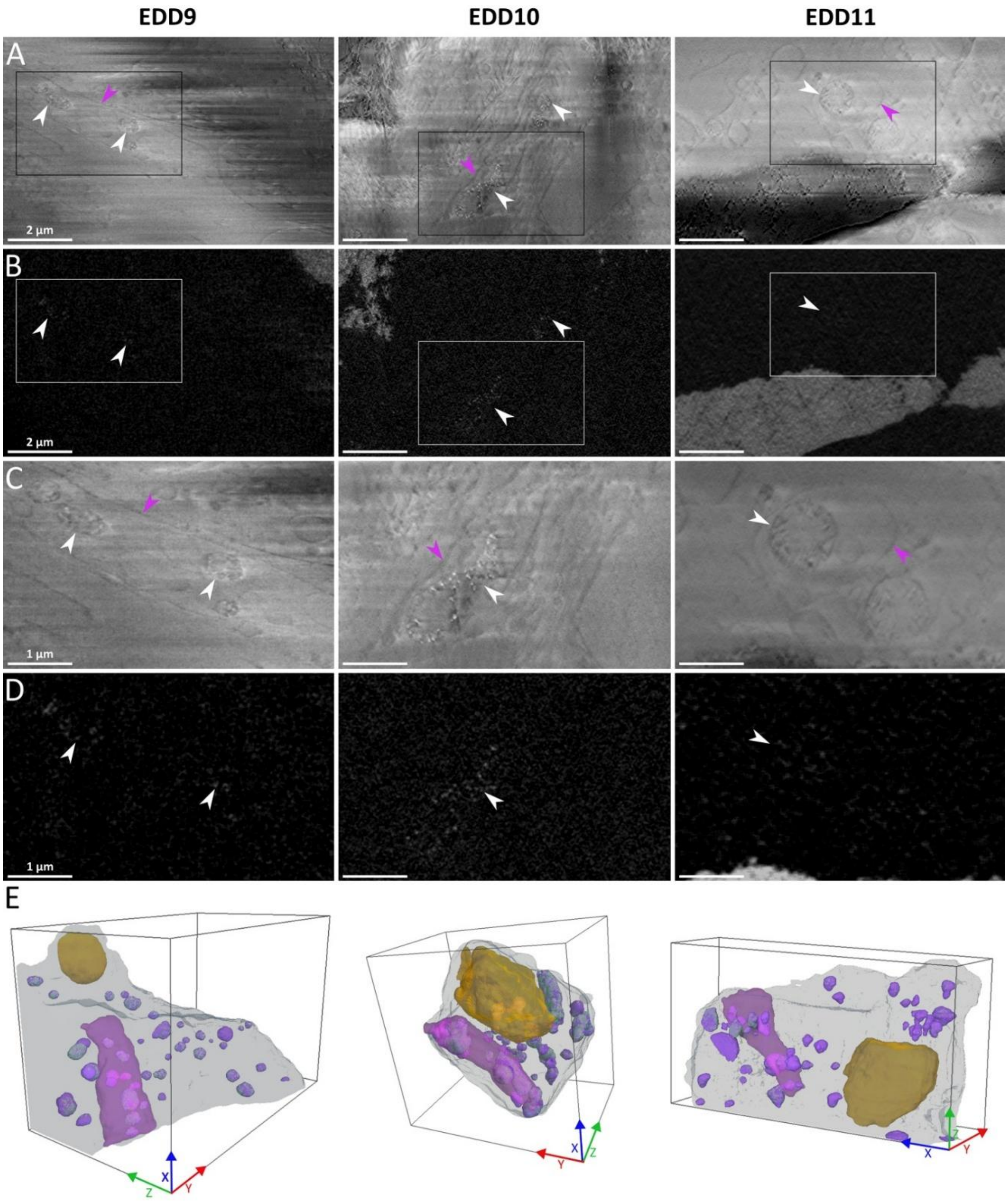


**Fig. S4.** Images of large membrane-bound structure containing vesicles filled with mineral precursors at EDD9 to 11 since they were not observed at EDD12 (A) Mixed InLens/SE image showing a large intracellular compartment delimited by a membrane (pink arrow) and containing multiple vesicles (white arrows) (B) Corresponding backscattered electron (BSE) image of the same region, revealing electron-dense particles within the internal vesicles (white arrows) (C) Three-dimensional perspective rendering illustrating the spatial organization of this membrane-bound compartment (pink) within the cell (light blue/grey) and the intracellular vesicles (purple) containing mineral precursors (green). Boxes dimensions : (EDD9) X 11.52 µm, Y 17.34 µm, Z 10.33 µm, (EDD10) X 11.37 µm, Y 11.59 µm, Z 8.11 µm and (EDD11) 22.73 µm, Y 6.10 µm, Z 12.09 µm.

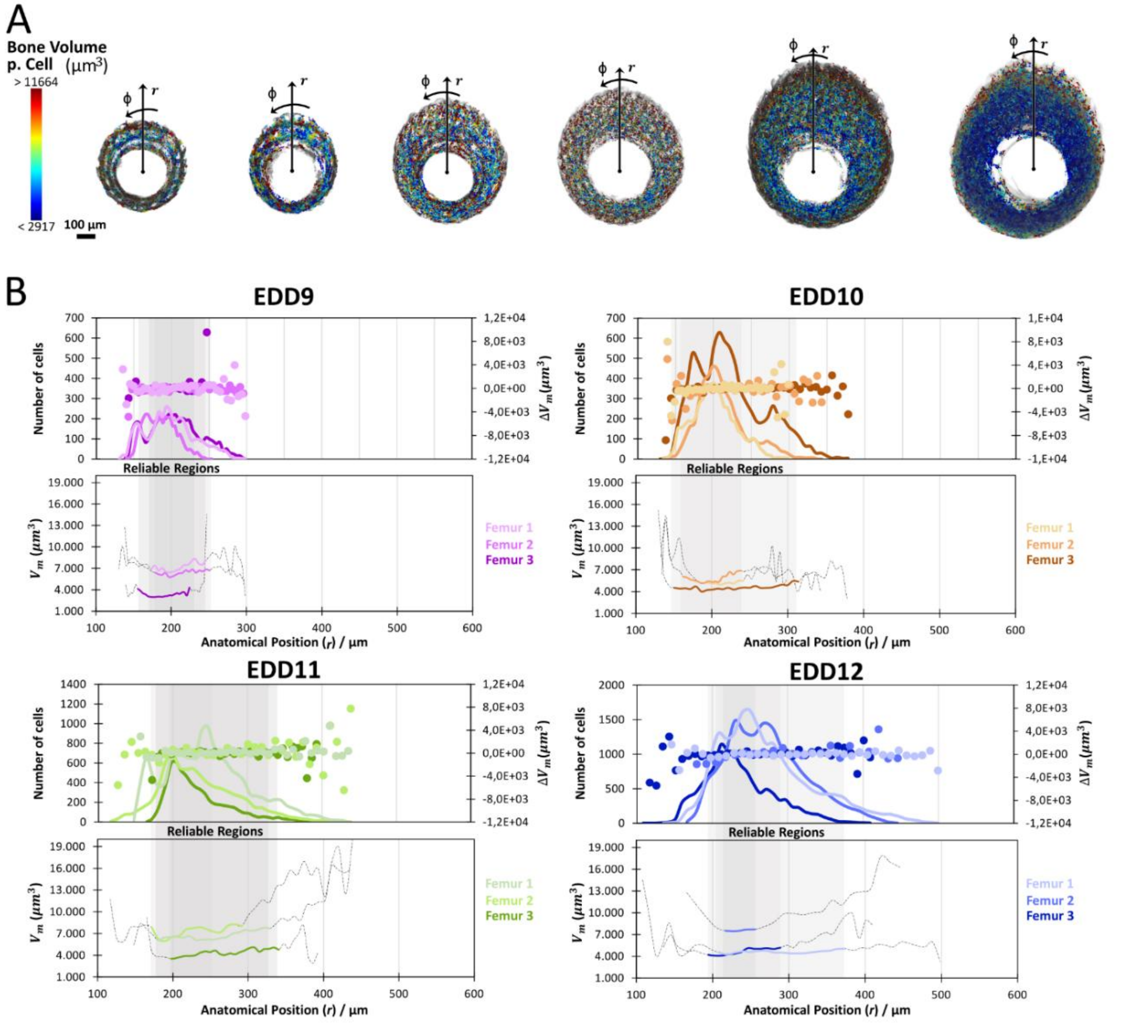


**Fig. S5.** Radial distribution of bone volume per cell at the femoral midshaft during development. A) Transverse cross-sections of representative femoral midshafts showing the spatial distribution of bone volume normalized per cell (µm³) in successive developmental stages. Each point represents the local bone volume associated with an individual cell, color-coded according to the scale shown. (r) correspond to the radial position and (φ) is the circumferential orientation. (B) Radial analysis of cellular distribution and bone volume per cell for EDD9 to EDD12. The graphs show the number of cells (points) as a function of anatomical radial position (r), together with the averaged bone volume per cell (Vm) of each radial local region. Shaded regions indicate the radial intervals used for quantitative analysis ("reliable regions"), selected to exclude endosteal and periosteal edge effects and to ensure consistent comparison between samples. Grey dashed lines represent the full radial profiles prior to region selection.

### Supporting Information Text

#### S6: Velocity of intracellular vesicles carrying mineral precursors

##### 1. Definition

To estimate the velocity $v$ of vesicles carrying mineral precursors within bone forming cells, we consider a 1D steady-state transport process. In steady state, vesicles are continuously produced and released to sustain the mineralization demand. The physical transport occurs along an intracellular path of length $d_0$, and the number of vesicles present in the cell is assumed to remain constant, according to the steady state assumption. The vesicle flow required to sustain mineralization is determined by the rate at which vesicles must be delivered to the mineralization site.

Here are the relevant definitions:

$v$ : intracellular vesicle velocity (µm/s), or its approximation $\theta$
$R_v$ : estimated vesicle release rate per cell (vesicles/s)
$V_m$ : measured mineralized volume per cell (µm$^3$)
$t$ : (unknown) time required to mineralize $V_m$
$V_{mp}$: average volume of mineral precursors contained in one vesicle ($\mu m^3$) inside a typical bone-forming cell
$V_{mpcell}$: estimated volume of mineral precursor contained in the vesicles inside a typical bone-forming cell
$T$: observation time, i.e. time between two consecutive measurements (one day)
$N_c$ : steady-state number of vesicles present in a typical bone-forming cell
$d_0$ : transport distance (µm)
$\rho$ : density of vesicles along the transport path (vesicles/µm)

##### 2. Transport distance $d_0$

The transport distance $d_0$ is defined as the maximal distance that a vesicle must travel from the source of calcium to the site of mineralization. The calcium source is assumed to be located at the trabecular surface (and correspond to the blood vessel), while the mineralization site is associated with the cellular domain surrounding an osteocyte. Accordingly, $d_0$ is estimated as the sum of half the trabecular thickness and half the average distance between two osteocytes:

$$d_0 = \frac{1}{2}\,(\text{trabecular thickness}) + \frac{1}{2}\,(\text{inter osteocyte distance})$$

This value represents a maximal transport path and is used as a conservative estimate of the intracellular transport distance. We should note that $d_0$ and $V_m$ are not independent quantities.

##### 3. Estimation of the vesicle release rate $R_v$

The vesicle release rate $R_v$ is determined by the mineralized volume $V_m$ that a cell is responsible for and by the time $t$ that it takes to complete the mineralization. Each cell is assumed to be responsible for mineralizing a given volume within a time scale $t$ with $t < T$, where $T$ is the observation time of one day. The volume $V_m$ is estimated using a Voronoi approach based on µCT measurements, which assigns to each cell the surrounding mineralized volume associated with its spatial domain. Since mineral precursors are transported within vesicles, the mineralization requirement can be translated into the number of vesicles that must be delivered per unit of time, based on the average mineral content $V_{mp}$ of a single vesicle. Thus, the vesicle release rate per cell is given by

$$\frac{V_m}{V_{mp} \cdot t},$$

which we approximate with the underestimation

$$R_v = \frac{V_m}{V_{mp} \cdot T},$$

where $T$ is the observation time of one day. In our calculations, we will express $T$ in seconds. Therefore, $R_v$ is the estimated vesicle release rate per second.

#### 4. The flow equation for the velocity

Based on the lower estimate of the rate $R_v$, we estimate a lower bound $\theta$ for the velocity $v$, by

$$\rho \cdot \theta = R_v$$

which leads to:

$$\theta = \frac{R_v}{\rho}$$

Thus, $\theta$ represents a lower bound for the vesicle velocity $v$ to sustain mineralization. To obtain a quantitative estimate of $\theta$, we assume that the vesicle density is given by:

$$\rho = \frac{N_c}{d_0},$$

where $N_c$ is the steady state number of vesicles present in a bone-forming cell. The idea behind this choice is that all vesicles are 'lined up' to be delivered to the site of bone formation according to the rate expressed by $R_v$. Then, by substituting $\rho$ into the equation for $\theta$, we have:

$$\theta = \frac{R_v}{N_c/d_0}$$

Which is equivalent to:

$$\theta = \frac{R_v \cdot d_0}{N_c}.$$

We can put everything together and after rearranging the terms we get the following formula for $\theta$:

$$\theta = \frac{V_m}{N_c V_{mp}} \frac{d_0}{T},$$

where we can recognize that $N_c V_{mp}$ is the total volume of mineral $V_{mpcell}$ in a bone-forming cell.

#### 5. Estimated value of the velocity

To estimate the velocity $\theta$ we start by making use of the last observation and we rewrite this equation as

$$\theta = \frac{V_m}{V_{mpcell}} \frac{d_0}{T},$$

where $V_{mpcell}$ is the amount of mineral precursor contained in the vesicles inside a typical bone-forming cell. The quantity $V_{mpcell}$ will therefore be expressed in $\mu m^3$.

#### 6. Linear Regression extrapolation for $V_{mpcell}$

From the FIB-SEM measurements, we have the amount of mineral volume contained in a large sample of cells together with the volume of the observed cells. We use this information to extrapolate, for each EDD and stack, per linear regression to obtain $V_{mpcell}$ for a cell of size 1000 $\mu m^3$. The reason for this extrapolation is that the cells observed in the FIB-SEM characterization are typically smaller than the target cell and therefore the amount of mineral precursor found in these cells is not the right quantity to estimate the velocity $\theta$. Extrapolating in $V_{mpcell}$ ensures that we will incorporate the correlation between number of vesicles per cell $N_c$ and the mineral volume per vesicle $V_{mp}$ in a typical bone-forming cell.

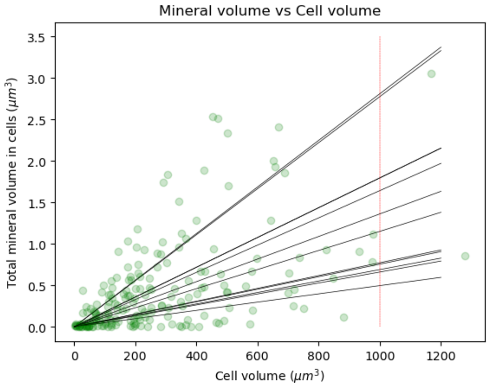


The dots indicate the volume of the single cells (x-axis) and the volume of mineral contained in the vesicles (y-axis). The 12 black lines are the linear regression lines. Their intersection with the red vertical line gives the volume of mineral content $V_{mpcell}$ in cells of size 1000 $\mu m^3$. The extrapolated values of $V_{mpcell}$ are given in the Jupyter notebook (Vesicle_Veleocity_V8).

### 7. Statistical analysis

We aim at determining if the velocity $\theta$ changes across the egg development days EDD.
To this purpose, for each of four EDDs (from EDD9 to EDD12), we have calculated the volume $V_m$ and we have computed the distance $d_0$, by µCT on three biological samples per development day. Therefore, the quantities $V_m$ and $d_0$ are not independent.
Furthermore, to determine $V_{mpcell}$ for each EDD we have characterized cells in three biological samples with FIB-SEM and used those cells to extrapolate for $V_{mpcell}$ at cell size of 1000 $\mu m^3$ for each sample.
Therefore, for each EDD we have six (independent) biological samples: three for µCT and three for FIB-SEM. For this reason, we will treat the quantities $(V_m \cdot d_0)$ and $V_{mpcell}$as independent quantities.
As mentioned, we evaluate the velocities defined from

$$\theta = \frac{V_m \cdot d_0}{V_{mpcell}} \frac{1}{T},$$

and consider the numerator and the denominator of the first fraction as two independent variables.
To simplify the notation, we will use the following variable names

$$X_{ij} = (V_m \cdot d_0)_{ij},$$

is the numerator, for EDD=i and Stack=j, with $i = 9,10,11,12$ and $j = 1,2,3$. Whereas

$$Y_{ik} = \left(V_{mpcell}\right)_{ik},$$

is the denominator, for EDD=i and Stack=k, with $i = 9,10,11,12$ and $k = 1,2,3$. Note that to avoid confusion, in this part of the calculation we are labeling the stacks differently to avoid confusion because, as mentioned earlier, the stacks for the microCT and the stacks for the FIB-SEM are independent.

**Detrended variables**
We define the detrended variables

$$\tilde{X}_{ij} = X_{ij} - \bar{X}_i,$$

and

$$\tilde{Y}_{ik} = Y_{ik} - \bar{Y}_i,$$

where $\bar{X}_i = \sum_{j=1}^{3} X_{ij}/3$ and $\bar{Y}_i = \sum_{k=1}^{3} Y_{ik}/3$ are the averages per EDD.

**Normality check**

We check if the detrended variables are normally distributed by using a q-q-plot for each of them. For $\tilde{X}_{ij}$ we have:

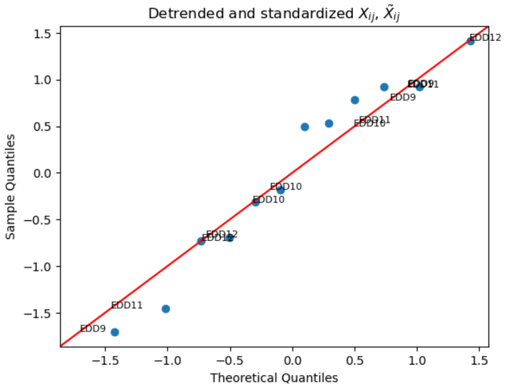


For $\tilde{Y}_{ik}$ we have:

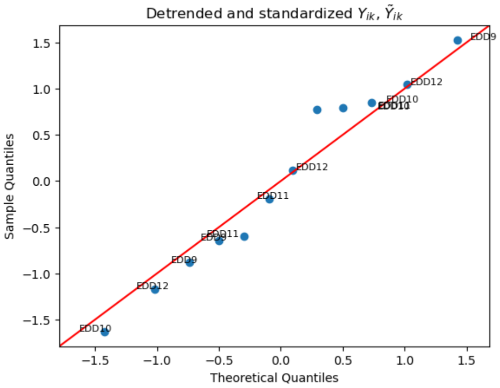


The above q-q-plots indicate that a large part of the data follows a normal distribution with the same variance. The detrended data was standardized for the purpose of the q-q-plot. We have also checked if any of the data is strongly suspicious to be an outlier. From the following plot we see that all datapoints are within 2*std thereby indicating that there are no further analyses needed:

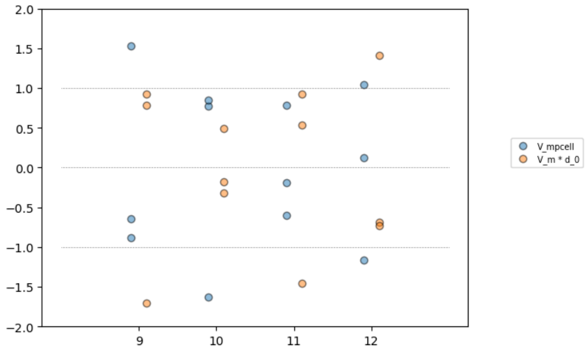


Nevertheless, we have 67% of the data in $V_{mpcell}$ within one std, and 75% of the data in $(V_m \cdot d_0)$ within one std. This may indicate that $(V_m \cdot d_0)$ has a distribution with thinner tails than normal. Nevertheless, we still conclude in this part of the analysis that considering the q-q-plots it is a good approximation to assume the detrended variables to follow a normal distribution. The variances associated to the detrended variables will then be computed using all available data.

**Standard errors for $\overline{X}_i$ and for $\overline{Y}_i$**

We start by estimating the standard deviation of the detrended $\tilde{X}_{ij}$ as

$$s_X = \sqrt{\frac{1}{11}\sum_{i=9}^{12}\sum_{j=1}^{3}\tilde{X}_{ij}^2}$$

and we associate to each $\bar{X}_i$ the same standard error

$$se_{\bar{X}} = \frac{s_X}{\sqrt{3}},$$

where the $\sqrt{3}$ arises from the fact that each $\bar{X}_i$ is the mean out of three independent microCT measurements.

In a very similar fashion, we continue by estimating the standard deviation of the detrended $\tilde{Y}_{ik}$ as

$$s_Y = \sqrt{\frac{1}{11}\sum_{i=9}^{12}\sum_{k=1}^{3}\tilde{Y}_{ik}^2}$$

and we associate to each $\bar{Y}_i$ the same standard error

$$se_{\bar{Y}} = \frac{s_Y}{\sqrt{3}},$$

where the $\sqrt{3}$ arises from the fact that each $\bar{Y}_i$ is the mean out of three independent FIB-SEM measurements.

**Theta and the standard error associated to theta**

For each EDD $i = 9,10,11,12$ we define the velocity of vesicles as

$$\theta_i = \frac{\bar{X}_i}{\bar{Y}_i}\frac{1}{T}$$

and with the use of the method of the propagation of the errors, we define the standard error associated to each $\theta_i$ from

$$\theta_i \pm se_{\theta_i} = \frac{\bar{X}_i \pm se_{\bar{X}}}{\bar{Y}_i \mp se_{\bar{Y}}}\frac{1}{T}$$

and expanding up to the first order

$$\theta_i \pm se_{\theta_i} \approx \frac{\bar{X}_i}{\bar{Y}_i}\frac{1}{T}\left(1 \pm \left(\frac{se_{\bar{X}}}{\bar{X}_i} + \frac{se_{\bar{Y}}}{\bar{Y}_i}\right)\right)$$

thereby leading to

$$se_{\theta_i} = \theta_i\left(\frac{se_{\bar{X}}}{\bar{X}_i} + \frac{se_{\bar{Y}}}{\bar{Y}_i}\right).$$

The values for the standard errors and for all variables rounded to the first decimal point are given in the following table

| EDD | Y_bar_i | se_Y | X_bar_i | se_X | theta | se_theta |
|---|---|---|---|---|---|---|
| 9 | 1.1 | 0.3 | 62191.7 | 10823.3 | 0.7 | 0.3 |
| 10 | 2.4 | 0.3 | 60846.0 | 10823.3 | 0.3 | 0.1 |
| 11 | 0.8 | 0.3 | 70915.3 | 10823.3 | 1.1 | 0.5 |
| 12 | 1.3 | 0.3 | 64337.7 | 10823.3 | 0.6 | 0.2 |

Note that the standard error associated to each $\bar{X}_i$ is the same and that also the standard error associated to each $\bar{Y}_i$ is the same because they have been computed using all detrended variables. The resulting values for the velocity are represented also in Figure 1 in the manuscript and reported here for convenience:

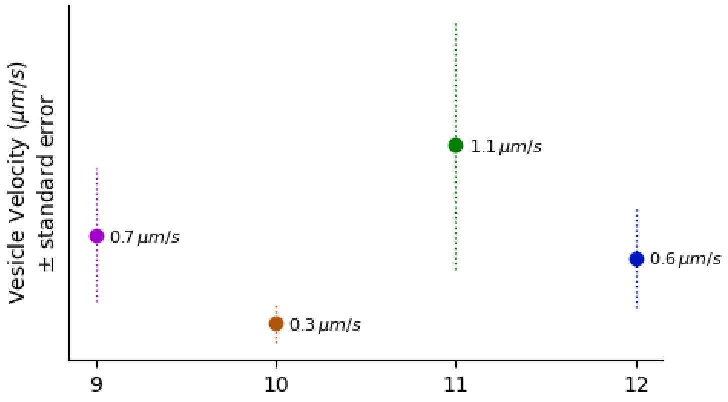


**Statistical significance**

We have compared the velocities theta pairwise by using a two-sided z-test with alpha = 0.1 as significance threshold. The statistic for each comparison between EDD=a and EDD=b is given by

$$stat_{ab} = \frac{|\theta_a - \theta_b|}{s_{ab}},$$

where

$$s_{ab} = \sqrt{(se_{\theta_a})^2 + (se_{\theta_b})^2}$$

and provided in the following table (rounded to two decimal points)

| EDD | **10** | **11** | **12** |
|---|---|---|---|
| **9** | 1.27 | 0.64 | 0.27 |
| **10** | | 1.42 | 1.21 |
| **11** | | | 0.85 |

The p-values are determined as

$$p_{ab} = 2\int_{stat_{ab}}^{\infty} \phi(x)dx\,,$$

where $\phi(x)$ is the standard normal distribution. The table of all p-values in the pairwise comparisons is provided in the following table (rounded to one decimal point)

| EDD | **10** | **11** | **12** |
|---|---|---|---|
| **9** | 0.2 | 0.5 | 0.8 |
| **10** | | 0.2 | 0.2 |
| **11** | | | 0.4 |

which indicate that no one of the pairwise comparisons is significantly different.

**Power analysis**

The missing significance of the pairwise comparison can be a reflex of the absence of differences or a consequence that the sample sizes are too small. We have therefore performed a power analysis to determine the power of the actual test under the assumption that the observed effect sizes are true.

We indicate the effect size of a pairwise comparison as

$$e_{ab} = |\theta_a - \theta_b|$$

and for $\alpha = 0.1$ determine $z_\alpha$ as

$$\alpha = \int_{z_\alpha}^{\infty} \phi(x)dx$$

where $\phi(x)$ is the standard normal distribution. Following the standard procedure, we compute the power $K$ as

$$K = \int_{z_\alpha - e_{ab}/s_{ab}}^{\infty} \phi(x)dx$$

thereby leading to the following values

| EDD | **10** | **11** | **12** |
|---|---|---|---|
| **9** | 0.5 | 0.3 | 0.2 |
| **10** | | 0.6 | 0.5 |
| **11** | | | 0.3 |

thus indicating that the missing significance may be due to lack of power. However, sample size calculations indicate that we would need more than 7 embryos per experimental technique in order to achieve a power of 0.8 in the comparison between EDD=10 and EDD=11 under the assumption that the effect size is as computed here.

**Permutation test**
For every two EDD, say EDD=a and EDD=b we assume as null hypothesis that the velocities are the same. To test against this hypothesis, we first consider the vectors $X_{aj}$, $X_{bj}$, $Y_{ak}$, $Y_{bk}$. We consider then the vectors

$$\boldsymbol{X} = (\boldsymbol{X}_a, \boldsymbol{X}_b)$$

And

$$\boldsymbol{Y} = (\boldsymbol{Y}_a, \boldsymbol{Y}_b)$$

each of which containing six elements, and we randomly permute the elements of these two vectors independently of each other.

**At each random permutation**, we therefore create two new vectors $\boldsymbol{X}^{(p)}$ and $\boldsymbol{Y}^{(p)}$ and take the first three elements of each for EDD=a and the second three elements for the EDD=b, thereby creating the vectors $\boldsymbol{X}_a^{(p)}$, $\boldsymbol{X}_b^{(p)}$, $\boldsymbol{Y}_a^{(p)}$, $\boldsymbol{Y}_b^{(p)}$.

We then compute the velocities $\theta_a^{(p)}$ , $\theta_b^{(p)}$ from the permuted variables and also the errors associated to them $se_{\theta_a}$ and $se_{\theta_b}$, taking into consideration that the standard errors associated to the $X$ and $Y$ variables are the same as computed above, because they were computed using all variables across all EDD.

Finally, we compute the statistic

$$stat_{ab}^{(p)} = \frac{\left|\theta_a^{(p)} - \theta_b^{(p)}\right|}{\sqrt{{se_{\theta_a}}^2 + {se_{\theta_b}}^2}}.$$

**By repeating this procedure** several thousands of times, we can compute the p-value as the proportion of $stat_{ab}^{(p)}$ that are larger than the true statistic. This leads to the following table of p-values:

| EDD | **10** | **11** | **12** |
|---|---|---|---|
| **9** | 0.2 | 0.4 | 0.7 |
| **10** | | 0.1 | 0.1 |
| **11** | | | 0.2 |

which are in the same level as the p-values computed with the z-test.
